# The effect of pH alterations on TDP-43 in a cellular model of amyotrophic lateral sclerosis

**DOI:** 10.1101/2022.12.21.521010

**Authors:** Yara Al Ojaimi, Charlotte Slek, Samira Osman, Hugo Alarcan, Sylviane Marouillat, Philippe Corcia, Patrick Vourc’h, Débora Lanznaster, Hélène Blasco

## Abstract

Amyotrophic Lateral Sclerosis (ALS) is the most common neurodegenerative disease affecting motor neurons. The pathophysiology of ALS is not well understood but TDP-43 proteinopathy (aggregation and mislocalization) is one of the major phenomena described. Several factors can influence TDP-43 behavior such as mild pH alterations that can induce conformational changes in recombinant TDP-43, increasing its propensity to aggregate. However to our knowledge, no studies have been conducted yet in a cellular setting, in the context of ALS. We therefore tested the effect of cellular pH alterations on the localization and aggregation of TDP-43. HEK293T cells overexpressing wildtype TDP-43 were incubated for 1h with solutions of different pH (6.4, 7.2, and 8). Incubation of cells for 1h in solutions of pH 6.4 and 8 led to an increase in TDP-43-positive puncta, an effect that was lost after 2h of incubation. This was accompanied by the mislocalization of TDP-43 from the nucleus to the cytoplasm. Our resulats suggest that small alterations in cellular pH affect TDP-43 and increase its mislocalization into cytoplasmic TDP-43-positive puncta, which might suggest a role of TDP-43 in the response of cells to pH alterations.

## Introduction

Amyotrophic lateral sclerosis (ALS) is a neurodegenerative disease characterized by the progressive death of motor neurons. This disease leads to the paralysis of skeletal muscles and ultimately respiratory muscles, resulting in the death of patients within 3 to 5 years following the onset of symptoms [1]. Abnormal aggregation of various proteins, particularly the TAR DNA binding protein 43 (TDP-43) has been reported in motor neurons and glial cells of ALS patients.

TDP-43 is a heterogenous nuclear ribonucleoprotein (hnRNP) that plays an essential role in transcriptional and post-translational regulation [2]. Physiological TDP-43 primarily acts as a nuclear protein, that shuttles between the nucleus and the cytoplasm. However, in pathological conditions, TDP-43 mislocalizes to the cytoplasm where it undergoes post-translational modifications and aggregation. These modifications have been associated with toxic loss and gain of function of this protein, leading to homeostatic imbalances.

pH changes have been described to have significant implications on the pathophysiology of neurodegenerative disorders [3]. Pathological mechanisms observed in ALS, such as ischemia, mitochondrial alterations, glutamate-mediated excitotoxicity and oxidative stress, all of which can affect intracellular pH, have been associated with the cytoplasmic mislocalization and aggregation of TDP-43 [4–6]. Concomitantly, several studies have shown that alterations in the pH can affect the structure of purified recombinant TDP-43 [7,8]. However, no studies have assessed the effect of such pH alterations on cellular TDP-43 in the context of ALS. The goal of our study was to assess the impact of pH alterations on the localization and solubility of cellular TDP-43, which emphasizes the importance of pH control when considering therapeutic strategies.

## Results

### pH-dependent cell viability

We first assessed the effect of extracellular pH changes on the viability of cells in order to determine the right incubation time. The incubation of HEK293T cells with different pH conditions for 1h does not decrease the viability of cells (**figure 1.A**). On the other hand, 4h and 24h incubation reduces the viability of the cells by more than 50% (results not shown). Therefore, the timepoint used in this study was limited to 1 hour of incubation. The viability of HEK293T cells transfected with 3 μg of plasmid expressing either GFP, TDP-43-6*His (TH), or GFP-TDP-43-6*His (GTH) and incubated with the different pH solutions for 1 hour was also assessed. No change in cellular viability was observed under these different conditions (**Figure 1 B-D)**.

**Figure 1:**
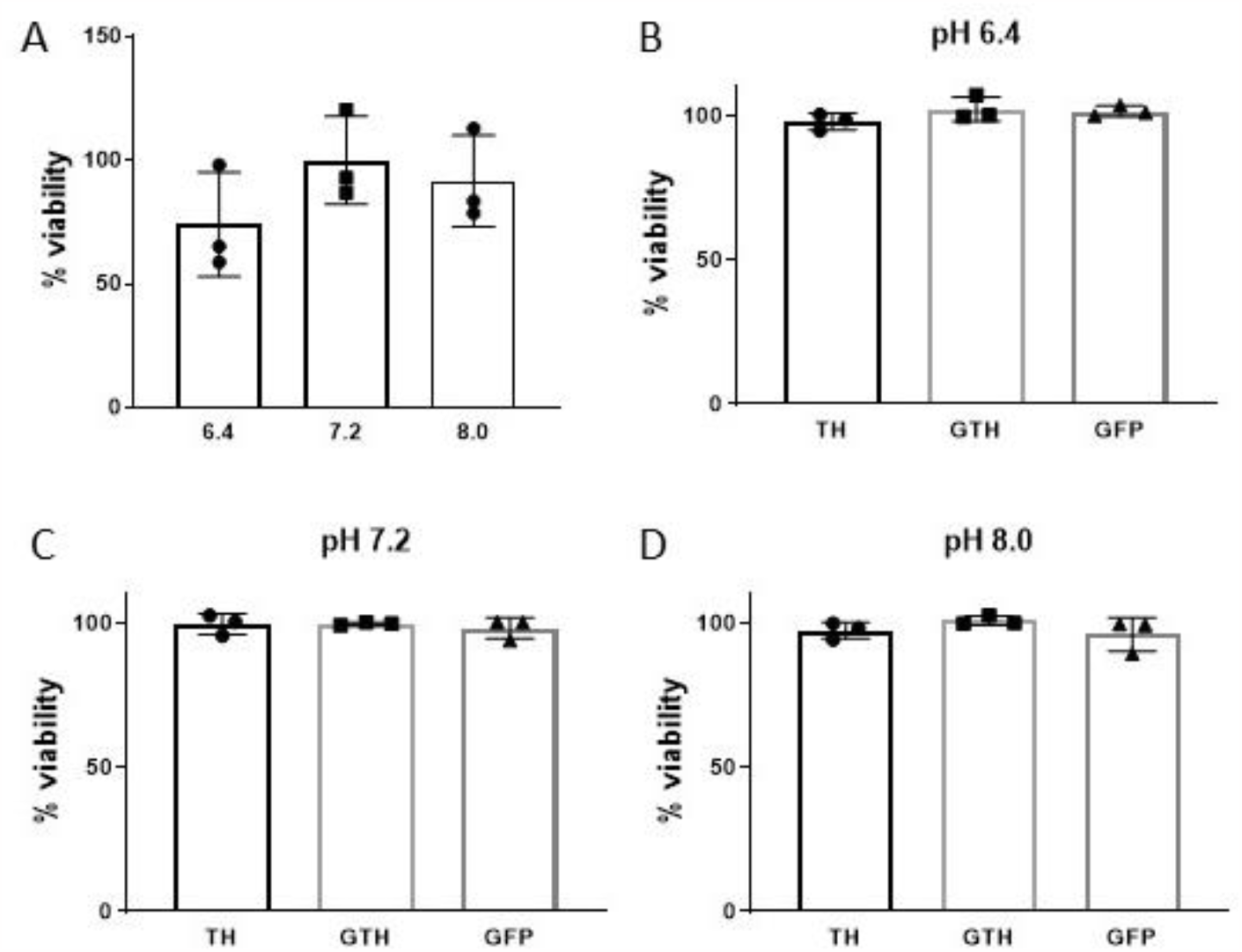
The viability of HEK293T cells is not decreased under different pH conditions **A)** MTT reduction assay of untransfected HEK293T cells incubated for 1h with different pH conditions. Values are expressed as percentage viability compared to pH 7.2 condition. Kruskall-Wallis test (N=3) **B-D)** MTT reduction assay of HEK293T cells transfected with 3 μg of plasmid expressing TDP-43-6*His (TH), GFP-TDP-43-6*His (GTH) or GFP and incubated for 1 hour at pH 6.4, 7.2 or 8. Values are expressed as percentage of viability compared to control (empty vector). Kruskal-Wallis test (N=3).

### pH-dependent TDP-43 localization

We then examined the effect of pH alterations on the localization of overexpressed TDP-43. At pH 7.2, most cells had TDP-43 mainly localized in the nucleus, with a small percentage of cells showing both nuclear and cytoplasmic TDP-43. Following a 1h incubation with solutions at pH 6.4 or pH 8, we detected an increase of around 6% in cells showing TDP-43 mislocalized to the cytoplasm compared to the control condition at pH 7.2 (**Figure 2. A,B**). We also observed an increase in the % of cells showing TDP-43-positive puncta at pH 6.4 (15.3 ± 1.8 %) and pH 8 (13.8 ± 0.9 %) compared to pH 7.2 (**Figure 2. A,C**).

**Figure 2.**
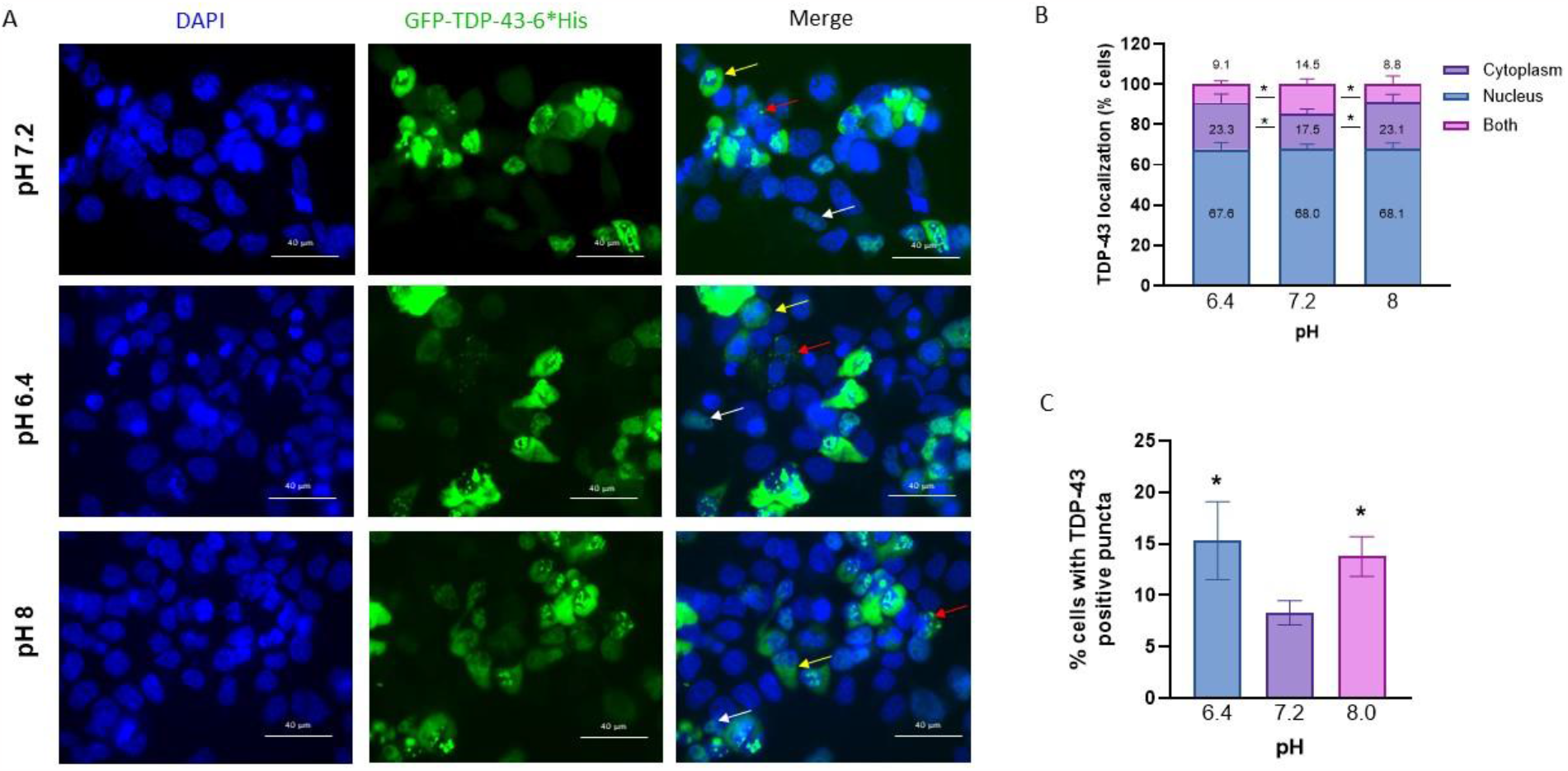
Changes in intracellular pH affect the localization of TDP-43 **A)** Immunofluorescence of HEK293T cells overexpressing GFP-wtTDP-43-6*His, following a 1h incubation at pH 7.2, pH 6.4, or pH 8. White arrow: nuclear TDP-43; Yellow arrow: cytoplasmic TDP-43. Red arrow: TDP-43-positive puncta. Scale bar = 40 μm **B)** Percentage of cells with TDP-43-positive puncta. \**p* = 0.04, Kruskal-Wallis test, N=4. **C)** Localization of TDP-43 under different pH conditions. \**p* = 0.04, two-way ANOVA, N=4.

### pH-dependent TDP-43 aggregation

The solubility of TDP-43 was also assessed by western blot. There was no change in the solubility of GFP under different pH conditions, indicating that the GFP tag has no effect on the solubility of TDP-43 under these conditions. Concomitantly, there was no difference in the solubility of TDP-43 when cells were overexpressing GTH compared to TH, confirming the lack of effect of GFP on TDP-43 solubility. We detected no change in the amount of insoluble full length TDP-43 **(Figure 3. A,B)** or insoluble 35 kDa C-terminal fragment of TDP-43 **(Figure 3. A,C)** at pH 6.4 or pH 8 compared to pH 7.2.

**Figure 3:**
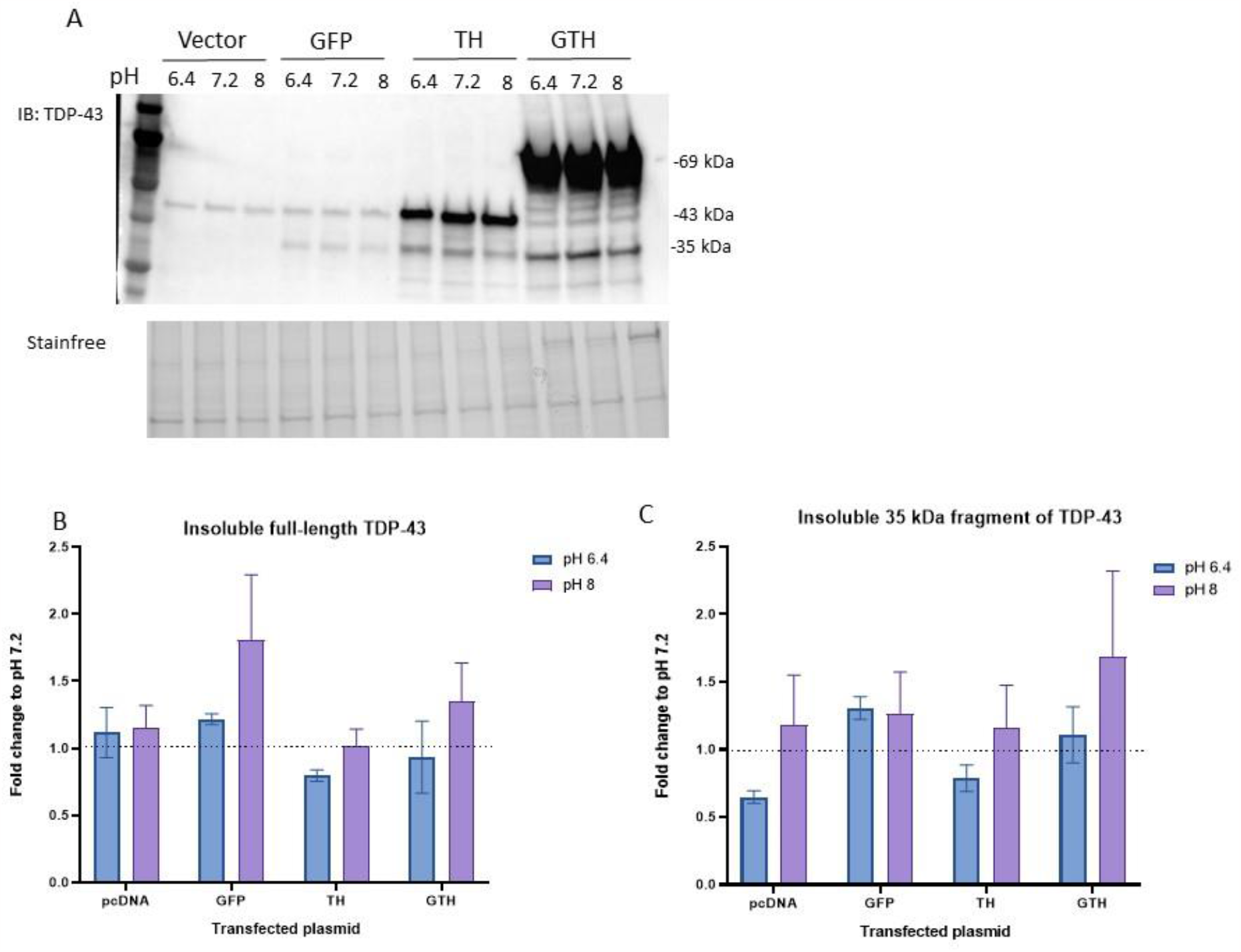
No change in the insolubility of TDP-43 following a 1h incubation with different pH conditions. Immunoblot **(A)** and quantification of insoluble full-length **(B)** and 35 kDa fragment **(C)** of TDP-43 in HEK293T cells transfected with 3 μg of empty vector or plasmid expressing GFP, TDP-43-6*His (TH), GFP-TDP-43-6*His (GTH) and incubated for 1 hour with different pH conditions. Values were normalized to the corresponding pH 7.2 condition. Wilcoxon test, N=3.

## Discussion

Extracellular pH alterations have been shown to induce parallel but less pronounced changes in the intracellular pH [10–13]. The change in intracellular pH was verified in our cells using the pH-sensitive BCECF (data not shown). How intracellular pH induces TDP-43 mislocalization and aggregation has not been well investigated. Since TDP-43 is known for its ability to spontaneously aggregate, particularly through its intrinsically disordered C-terminal domain (CTD) [13], [14], it is complicated to assess the effect of microenvironmental alterations on the full-length protein. Therefore, most studies that investigated the effect of pH on TDP-43 folding have used isolated protein domains rather than the full-length protein [15– 17]. In addition, the results obtained seem to vary according to the recombinant TDP-43 forms used and the protocols applied (i.e. protein concentration, temperature, buffer pH, salt concentration). Although several studies show that TDP-43 domains can be structurally altered upon pH changes [18–21], the impact observed on separate protein domains might not be reproducible in the full-length protein given the presence of complex intra- and inter-molecular interactions [22], [23]. In addition, all these studies were performed on recombinant purified proteins, and not in cellular models of TDP-43 proteinopathy. Given the extensive post-translational modifications that TDP-43 can undergo inside cells [23], such studies do not take into consideration these modifications that can highly affect the structure and solubility of proteins.

Here we showed, for the first time, that pH alterations in the cellular environment can affect the cytoplasmic mislocalization of TDP-43. In cells overexpressing TDP-43, we observed a decrease in the nuclear localization of TDP-43 and an increase in the number of cells with TDP-43-positive puncta at pH 6.4 and 8 compared to the physiological pH. Western blot analysis revealed no significant change in the amount of insoluble TDP-43 at pH 6.4 and pH 8 in HEK293T cells overexpressing TDP-43. This discrepancy in the results observed through western blot and those observed through fluorescence microscopy can be explained by the differing accuracy of these two methods in detecting real insoluble aggregates. In fact, Barmada and his colleagues demonstrated that the TDP-43-positive puncta observed by immunofluorescence can sometimes be removed by detergent treatment, indicating that they can’t always be considered as insoluble TDP-43 aggregates [24]. These results suggest that studying TDP-43 aggregation by immunofluorescence can lead to its overestimation and that western blot analysis might be a more accurate method.

TDP-43 is particularly sensitive to changes in its environment which can shift the equilibrium towards favouring the formation of oligomeric forms [7]. Under stressful cellular conditions, TDP-43 undergoes liquid-liquid phase separation (LLPS) and is recruited to stress granules [24,25]. For instance, stress granules can be formed in the context of metabolic dysfunction caused by starvation, hypoxia, or mitochondrial abnormalities [26–30], which can enhance TDP-43 LLPS and subsequent aggregation. For instance, a recent study has shown that the regulation of the oxidative status of TDP-43 particularly at the level of the CTD, which is largely affected by the metabolic status of the cell, affects the conformation of TDP-43 and its self-assembly [32]. Metabolic stressors are partly linked to intracellular pH changes and can affect the protonation state of charged amino acid residues and therefore alter the structure and conformation of proteins [31, 33]. Such modifications might allow proteins like TDP-43 to act as biosensors of alterations in the intracellular environment [34]. Our results therefore suggest that an acute change in intracellular pH is associated with increased mislocalization of TDP-43 and its assembly into cytoplasmic puncta, possibly as part of the cellular response to the pH stress. This highlights the importance of controlling for cellular pH when assessing potential therapeutic approaches in ALS. In addition, different cell types react differently to metabolic alterations and pH changes, with more or less efficient buffering mechanisms. Therapeutic approaches should therefore be designed accordingly and consider restoring cellular homeostasis in a cell-specific way. Future studies should give more focus to the effect of pH alterations on TDP-43 in neurons, which are particularly sensitive to changes in their environment, and assess the effect of regulating cellular pH on the proteinopathy associated with TDP-43.

## Materials and methods

### Cells and transfection

HEK293T cells (ATCC, USA) were cultured in Dulbecco’s modified Eagle’s medium (DMEM) supplemented with 10% (*v/v*) fetal bovine serum and 1% non-essential amino acids at 37°C and 5% CO_2_ and plated in six-well plates at a concentration of 0.3 × 10^6^ cells/well the day before transfection.

Cells were transfected at 60-70% confluency using the JetPEI transfection reagent (Polyplus transfection) following the manufacturer’s protocol. Briefly, a volume of JetPEI equivalent to twice the amount (μg) of the plasmid was used. The plasmid and JetPEI were prepared in sterile 150 mM NaCl. Following gentle mixing and an incubation for 15 min at RT, the mix was added drop by drop to the cells. All transfections used 3μg of plasmids unless otherwise mentioned, and lasted 48h.

### pH solutions

pH solutions were prepared to induce changes in intracellular pH, as previously described [35]. The stock solution was prepared with 125 mM NaCl, 4.5 mM KCl, 20 mM Hepes, 20 mM MES, 1 mM CaCl_2_ and 1 mM MgCl_2_. Once this solution was obtained, the pH was adjusted with NaOH or HCl in order to obtain two different solutions of pH 6.4, 7.2, and 8. Cells were incubated with these solutions for 1h before different assays were performed to determine the effect of pH changes on cell viability and TDP-43 localization and aggregation.

### Cell viability

Cell viability was examined using the MTT reduction assay. 48 h after transfection, and following incubation with the different pH solutions, the cell medium was removed, and cells were incubated with 0.5 mg/mL MTT for 30 min at 37°C. The MTT solution was then removed and DMSO was added to the cells. The plate was wrapped in foil and put on an orbital shaker for 10 min. Absorbance was then measured at 570 nm using a microplate reader (Biorad).

### Assessment of TDP-43 localization and aggregation

To study the effect of intracellular acidification on the localization of TDP-43, HEK293T cells were plated on coverslips pre-treated with poly-D-lysine (10 ug/mL) and transfected with the plasmid expressing GFP-wtTDP-43-6*His (3 μg, GTH). 48 h post-transfection, cells were incubated for 1 h with the different pH solutions (6.4, 7.2 and 8) as previously described. Cells were then incubated with 1μL of Hoechst 33342 (1:1000) for 10 min before reading the fluorescence. Afterwards, cells were fixed for 20 min with 4% paraformaldehyde solution (diluted in PBS). Images were obtained by confocal microscopy (Leica SP8) where 100 cells per condition were analysed for TDP-43 localization (nuclear, cytoplasmic, or both) and aggregation.

Western blot was used to assess the solubility of TDP-43. HEK293T cells were transiently transfected with 3 μg of plasmids coding for TDP-43-6*His (TH), or 3 μg of plasmids coding for GFP-TDP-43-6*His (GTH). The empty pcDNA6.2 plasmid and a GFP-coding plasmid were used as control (3 μg). 48 h after transfection, cells were treated for 1h with the different pH solutions before protein extraction.

The medium of HEK293T cells was removed and the cells were scraped off in ice-cold PBS and centrifuged at 900 x g for 5 min at 4ºC. Cells were then resuspended in 100 μL of cold lysis buffer (PBS pH = 8.0, 1% Triton-X 100, 10 mM MgCl_2_, 1 mM DTT) supplemented with 1X Halt Protease and Phosphatase Inhibitor Cocktails (*Thermo Fisher Scientific*) and Pierce Universal Nuclease (*Thermo Fisher Scientific*). After 30 min, 50 μL of lysate was centrifuged at 13000 x g for 5 min. The pellet, which makes up the insoluble protein fraction, was resuspended in RIPA supplemented with 6M urea. 10 μg of proteins were migrated by SDS-PAGE, transferred onto a 0.2μm PVDF membrane, and blocked with 5% fat-free milk in TBS-Tween (0.1%) for 1 h at room temperature. Membranes were then probed overnight at 4°C with a primary antibody diluted in the blocking solution at 1:5000 targeting either the C-terminus of TDP-43 (Rabbit, 12892-1-AP, *Proteintech*). The following day, membranes were incubated for 1 h at room temperature with a complementary HRP-conjugated secondary antibody diluted at 1:10000 in the blocking solution. Detection of proteins was achieved following 5 min incubation with Enhanced Chemiluminescence (ECL) reagent and exposed using a ChemiDoc imaging system (BioRad). Loading control was performed by normalizing the target protein to the stain-free images of total protein. All bands were quantified using the BioRad Image Lab software version 6.1.0.

### Statistical analysis

All statistical analyses were carried out using the GraphPad Prism 8 software. Data is reported as mean ± Standard Error of the Mean (SEM) for 3-5 independent experiments. The localization of the TDP-43 protein was analysed using a two-way ANOVA test. The influence of pH on TDP-43 puncta were analysed using the Kruskal-Wallis test. Analysis of the western blot results was done using the Wilcoxon test. Results were considered to be statistically significant at \**p* values < 0.05.

## Declarations

### Data availability

Data generated in this study is available upon reasonable request to the corresponding author.

## Acknowledgments

The authors would like to thank LabEx MAbImprove (ANR-10-LABX-53-01) and the Région Centre Val de Loire for their financial support.

## Authors’ contributions

Y.A., C.S, S.M., and D.L. performed the experiments and generated the data. Y.A., C.S, and D.L. analyzed the data. Y.A., C.S., and D.L. designed the experiments. Y.A., C.S., D.L. and H.B. wrote the manuscript. P.V. and S.O. revised the manuscript. All authors contributed to the article. All authors read and approved the final manuscript.

## Abbreviations

ALS: Amyotrophic lateral sclerosis
BCECF: 2’-7’-bis(carboxyethyl)-5(6)-carboxyfluorescein
CTD: C-terminal domain
HEK293T: Human embryonic kidney cells 293T
hnRNP: Heterogenous nuclear ribonucleoprotein
LLPS: Liquid-liquid phase separation
TDP-43: TAR DNA binding protein 43

